# Temporal dynamics of hippocampal activity predict stable patterns of sensitization or habituation to noxious stimulation across sessions

**DOI:** 10.1101/2023.09.06.555302

**Authors:** Richard Harrison, Carien M. van Reekum, Greig Adams, W. Gandhi, Tim V. Salomons

## Abstract

Acute pain serves to warn an organism of potential damage. When nociceptive stimulation persists, two possible responses emerge: If no risk of harm is anticipated, habituation may occur. If harm is considered possible, pain sensitization is likely. An individual’s adaptation to prolonged pain may provide insight into their ability to manage resources, and possibly their likelihood of developing chronic pain. Yet, little is known about the stability of these individual differences or their underlying neural mechanisms. Eighty-five participants undertook a repetitive noxious stimulation task and a resting-state scan in an MRI scanner, in a first session. They then completed the same task outside the scanner on three separate days. Pain adaptation was operationalized as the slope of change in pain ratings within session. Intraclass correlations were calculated between slopes across the four sessions, which demonstrated high stability and association with emotional disposition. Individuals who habituated to repeated stimuli showed increasing activity in the anterior hippocampus and amygdala, while individuals who sensitized showed increasing activity in the sensorimotor cortices. These clusters were then used as seeds in resting state analysis, with habituation associated with higher functional connectivity between hippocampus/amygdala and ventromedial prefrontal cortex(vmPFC), and higher connectivity between sensorimotor regions and the hippocampus, amygdala and insula cortex. Our findings suggest that pain adaptation is a stable phenotypic trait, which may have implications for the prediction of chronic pain.

This study implicates neural sensory and appraisal systems in these stable responses, offering insight into the mechanisms underlying trait-like responses to prolonged nociceptive input.

## Introduction

Acute pain serves as a warning of potential harm or damage. In response to repetitive and consistent nociceptive stimulation, two opposing adaptive responses are possible: Pain sensitization might occur if the source of persistent injury is perceived as a risk[74]. Alternatively, when experiencing repetitive stimuli that are unavoidable but non-harmful, the preservation of resources expended on defensive behaviour is adaptive, thus facilitating pain habituation[57].

In experimental contexts, it is frequently reported that individuals habituate to pain[13,22,25,27,29,37,50,68,70,73]. However, multiple studies demonstrate the opposite (sensitization)[19,22,32,45,51]. Most studies employ group-level analyses only, and simply report descriptives of the whole sample[8,36,59,73,75], or only include incomplete individual data[13,23,29,32,45,50,77]. Sample composition varies, from predominantly sensitizers[36] or habituators[29,68] to near equal splits[8,59,75]. This heterogeneity may merely represent differences in experimental paradigms, alternatively, adaptation profiles may vary within samples systematically. Findings suggest that the proportion of habituators in a sample may be stable over time[7], although these stem from a small study(n=10) and further investigation is required.

Pro- and anti-nociceptive modulatory mechanisms have been well documented[1,9,56,78], and phenotypic nociceptive profiles[85] are at the core of research investigating biomarkers of pain sensitivity and pain chronicity[21,30,54,86]. Many studies adopt single-session quantitative sensory testing (QST) techniques such as temporal summation (TS), to quantify pain sensitization. Psychological characteristics, such as depression, anxiety and neuroticism have also been identified as phenotypic profiles for chronic pain and treatment response[14,47,58,73,76]. To better understand these markers, profiles of pain adaptation need to be investigated over multiple sessions.

The neural underpinnings of acute pain perception involve a combination of higher-order cortical processes, alongside sensorimotor processing[55,56]. Prefrontal regions play an active role in pain modulation, with direct projections to opioidergic brainstem regions, such as the periaqueductal gray(PAG) and rostral ventromedial medulla(RVM)[16,62]. In addition, regions such as the cingulate and ventromedial prefrontal cortices(vmPFC) interact with subcortical brain regions, including the hippocampus and amygdala, to execute threat-based learning, update expectations, and appraise nociceptive stimulation[39,64,89], suggesting they may be involved in context-dependent adaptation of responses to nociceptive stimulation.

Repetitive stimulation has been associated with decreasing activity in sensory-discriminative regions, such as the somatosensory cortex, anterior insula and supplementary motor area[8,50,59], as well as increasing activity in medial prefrontal regions[7,8]. These may represent appraisal-based modulatory processes, alongside decreases in sensory-discriminative processing. However, these adaptation studies quantified habituation by comparing brain activation during nociceptive stimulation in early vs late MRI blocks[50,59] or across days of stimulation[8]. As pain ratings were not included in their analyses, these findings may simply represent adaptation to nociception, rather than to the subjective experience of pain.

This study aims to examine the stability of individual differences in pain adaptation to repetitive nociceptive stimulation across multiple days. We further test the extent to which these individual differences are reflected in engagement of distinct somatosensory and cortical networks, and with well-established psychometric and sensory pain indices. The combination of these approaches will enhance our understanding of phenotypical profiles of pain adaptation, and mechanisms that may underlie resilience or vulnerability to persistent pain.

## 2. Methods

### 2.1 Sample

We tested ninety-five participants between the ages of 18 and 45 years. Over the course of the experiment, 10 participants were excluded from analysis due to excessive head-movement during MRI data acquisition(n=2), malfunction of the thermal stimulator(n=7) and data corruption during data transfer(n=1). This, therefore, left a final sample of eighty-five participants (44 females, M_age_=23.45, SD=4.48y). Participants were recruited using opportunistic sampling via posters, email lists and social media advertisements across the University and local community. Participants were recruited to take part in 13-sessions, with the focus of this current study being on data from the first 5 sessions. This current study analysed a subset of the data from a larger study, for which the a-priori sample size of 90 participants was calculated, based on the delivery of a between-groups intervention.

All participants were right-handed, and reported no historical or current chronic pain diagnoses, neurological disorders or neuropathic conditions. Participants were asked to abstain from pain medication on the day of a session, were all right-handed and had no MRI counterindications. The study was approved by the ethics committee of the University of Reading (UREC14/04). All participants provided fully informed written consent and received reimbursement (£10/hour) for their participation.

### 2.2 Materials

#### 2.2.1. Equipment

Thermal nociceptive stimulation was administered using a Medoc Pathway ATS device[9]. The thermode was a 30×30mm Peltier thermode.

For use in the MRI, the Medoc Pathway system was fitted with an MR-filter, to reduce the influence of extraneous noise within the MR environment. Experimental stimuli in the scanner were delivered using E-Prime 3 (Psychology Software Tools, Pittsburgh, PA), where responses were provided via a fibre-optic, 4-button response pad.

#### 2.2.2. Questionnaires

To investigate the psychological features of pain adaptation profiles, participants completed the Big Five Inventory (BFI;[8]), Beck‘s Depression Inventory (BDI;[3]) and Five Factor Mindfulness Questionnaire (FFMQ;[2]), which were collected during sessions 3 (BFI & BDI) & 4 (FFMQ). The BFI is a 44-item inventory which captures extraversion (vs introversion), agreeableness (vs antagonism), conscientiousness (vs lack of direction), neuroticism (vs emotional stability) and openness (vs closedness to experience). The resulting score is subdivided across these five subscales. The BDI is a 21-item scale that measures symptoms of depression. A high score on the BDI represents higher symptoms of depression. Lastly, the FFMQ is a 39-item questionnaire based on a factor analysis of 5 independently developed mindfulness questionnaires. This measure provides a total score for trait mindfulness (higher scores represent higher mindfulness) across five subscales: openness, description, acceptance, non-judgementalness and non-reactivity. Within this sample, the BDI (α=.84) and FFMQ (α=.87) demonstrated high stability and consistency. This was also the case for each of the five subscales of the BFI, extraversion (α=.85), agreeableness (α=.74), consciousness (α=.83), neuroticism (α=.85) and openness (α=.75).

Pain ratings were provided on an 11-point numerical rating scale (NRS), ranging from 0(no pain) to 10(extremely painful). Participants were instructed that, regardless of whether they can feel heat or not, if they do not consider the stimulus to be painful, they should provide a 0. Participants were also informed that they could provide ratings in intervals of 0.5 along this scale.

#### 2.2.3. MRI Acquisition

Brain images were acquired using a 3-Tesla Siemens Magnetom Prisma (Siemens, Erlangen, Germany) and all images were acquired using a 64-channel head and neck coil. The narrow size of the coil restricted head movement, although in the instance of smaller head sizes, additional foam padding was used to restrict movement. The scanning protocol consisted of anatomical and functional imaging, all of which utilised an interleaved spatial order acquisition. A T1-weighted inversion recovery fast gradient echo-high resolution anatomical scan (TR=2.3s, TE=2.29ms, FA= 8°, voxel size=0.9x.0.9×0.9mm, 256×256×192 matrix), T2^*^-weighted gradient echo planar imaging(EPI) resting-state sequence(TR=2.29s, TE=36.4ms, FA=84°, volumes=210, multiband acceleration factor=2, voxel size= 2.1×2.1×2.1mm, slice thickness=0.94mm, matrix=84×84×58) and four EPI pain stimulation blocks(TR=2.29s, TE=36.4ms, FA=84°, volumes=168, multiband acceleration factor=2, voxel size=2.1×2.1×2.1mm, slice thickness=0.94mm, matrix=84×84×58) were collected for each participant. For the purposes of creating field maps, two spin-echo (SE) EPI scans in opposite encoding directions (PA;AP) were completed prior to the four pain EPI blocks.

### 2.3 Procedure

#### 2.3.1. Baseline Assessment (Session 1)

All participants attended an initial baseline session, wherein they were briefed on the study timeline, provided informed consent, and completed a quantitative sensory testing (QST) battery. Prior to the initiation of QST, participants were given an overview of the Medoc Pathway system (Medoc Medical Systems, Ramat Yishai, Israel), to reduce situational anxiety towards the equipment. Participants were also given a description of the NRS that they would be using to provide pain ratings when prompted. The thermode was placed in a custom-made wooden leg rest and placed next to the participant’s chair. Participants then placed the underside of their lower right calf over the stimulus.

Firstly, participants completed a pair of tasks to determine their pain threshold, as described in previous publications[31]. The resulting thresholds from 1) a staircase/method of levels experiment and 2) a method of limits experiment were averaged to calculate an overall participant pain threshold.

Participants then completed a percept calibration test, with a 20s ramp and hold stimulus, with ramp and return rates of 8°C, to reduce the risk of ceiling (intolerably painful) or floor (non-painful) effects. Based on the results of piloting, the destination temperature was set as threshold+0.5°C. After the stimulus, the participant provided a pain intensity rating using the 11-point NRS. If the participant rated the stimulus between 4-6, the temperature was formalised as their test temperature. If the participant rated outside of this range, the temperature was altered +/-0.5°C, and the test was repeated until a rating between 4-6 was provided.

The participant then completed experiments to quantify temporal summation (TS), as well as conditioned pain modulation and intrinsic attention to pain tasks (not reported here). For TS, a phasic design was used, which consisted of a 120s ramp and hold stimulus, using the calibrated test temperature, with ramp and return rates of 8°C. Participants were asked to provide pain intensity ratings, when prompted, at 10s intervals, ultimately providing a total of 12 ratings. TS was calculated as the difference between the first and last rating, with a positive value representing enhanced sensitization and higher TS.

Lastly, the stimulus for the habituation paradigm was calibrated, as a higher intensity of 6-8 was required. The calibration consisted of three 20s stimuli, with a destination to temperature of threshold + 1°C. Participants provided a pain intensity rating for each stimulus. If all three ratings were between 6-8, the habituation stimulus was set. If they rated outside of this range, the temperature was altered by +/-0.5°C, and the test was repeated until acceptable ratings were received.

#### 2.3.2. MRI Imaging Session (Session 2)

Participants attended an MRI session shortly after completing their baseline assessment (Mean=2.4 days, range= 1-11 days). Participants were instructed to keep their body and, specifically, their head still. They were also instructed to keep their eyes open and to focus on a fixation cross projected via a monitor and mirror attached to the head coil. After an initial localiser scan, structural images were acquired using a 5min 21s T1 anatomical scan, which was followed by an 8min 10sec resting-state scan. Participants then received the task instructions a second time and were given a test stimulus to their left leg, using the calibrated temperature from the baseline assessment, to check tolerability which all participants confirmed. The pain task (Fig.1) was divided across four EPI scans. Each block consisted of 11 stimuli, with a duration of 8s, ramp and return rates of 8°C and an interstimulus interval (ISI) of 20s. At the end of each block, participants were asked to provide an average pain rating of the preceding stimuli, using a 4-button response pad. Participants were instructed to tap the furthest left-hand button to move the rating slide to the left, and vice versa for the far-right button. The two spin-echo field maps were then collected to finish the scanning session.

**Figure 1.**
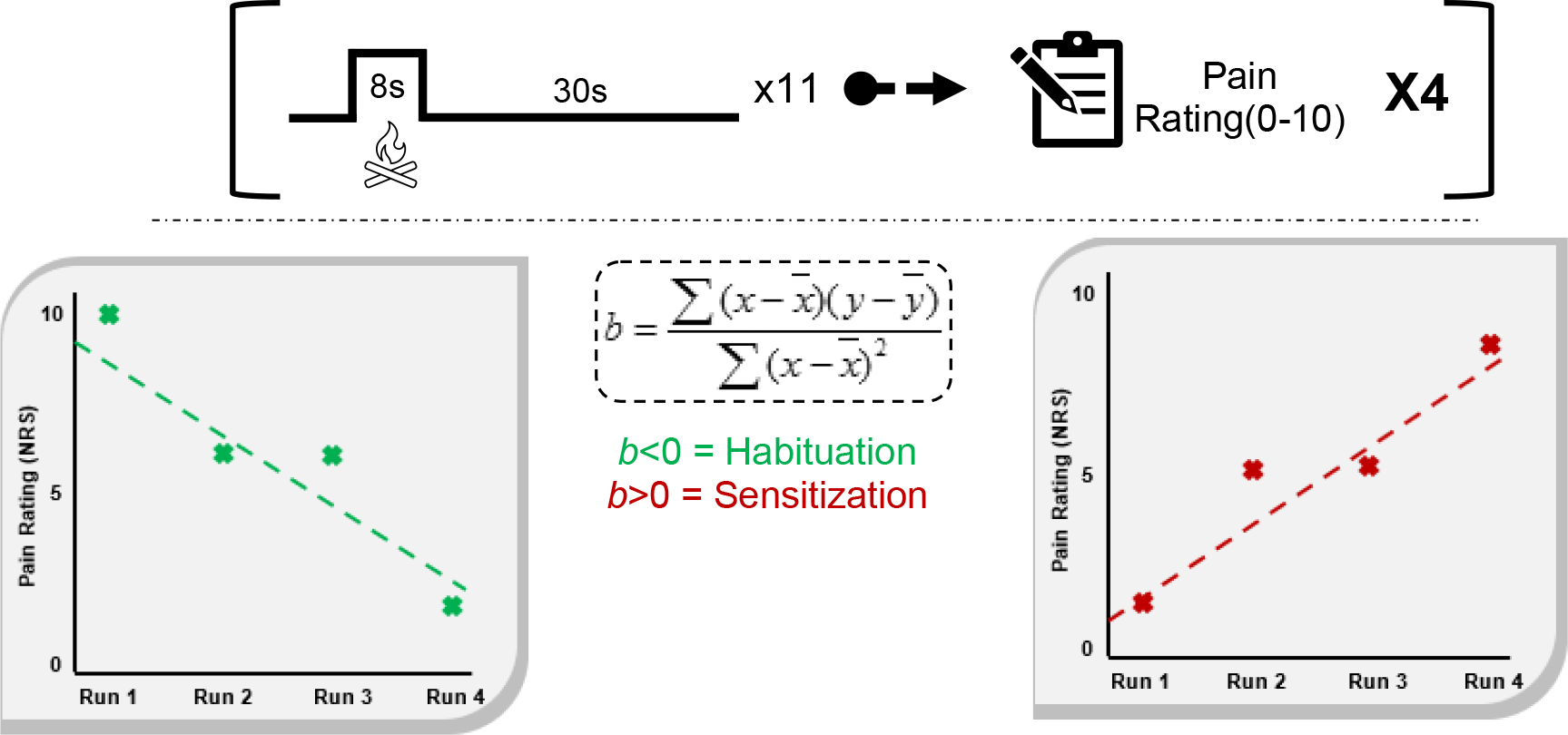
Experiment Design | Each block(top) consisted of 11 thermal stimuli. After the stimuli were administered, participants were asked to provide an average pain intensity rating for the stimuli that preceded. After four blocks, a linear regression equation was fitted to the points and an angle of slope was calculated. The quantification of pain adaptation(habituation/sensitization) was based on this slope, with a negative value representing habituation(bottom-left), and a positive value for sensitization(bottom-right). Data presented in these graphs is hypothetical for explanatory purposes, with block number on the x-axis, and with the provided pain rating for each block on the y-axis.

#### 2.3.3. Pain Stimulation Sessions (Sessions 3-5)

The following three sessions were completed following the imaging session (Mean=3.9 days, range= 1-9 days). These pain stimulation sessions consisted of psychometric assessment, wherein participants completed questionnaires at the beginning of each session. After this, participants completed the same task as they received in the MRI scanner, with the thermode applied to their left calf. As such, each participant received four blocks of a series of 11-stimuli, providing an average pain intensity rating for each train of stimuli.

### 2.4 Data Reduction and Analysis

#### 2.4.1 Behavioural Data Analysis

Previous studies have identified weakness of characterising habituators and sensitizers via dichotomous splits, as this approach forces participants with very little change into a distinct bipolar comparison[46,75]. As such, pain adaptation was quantified as a continuous variable by calculating the y-intercept of the regression slope across the pain ratings provided at the end of each of the four blocks of pain stimulation, administered during the MRI. The calculation of adaptation as a continuous variable meant that a negative value indicated habituation to pain over time, via decreasing pain ratings, whereas a positive value represented sensitization (Fig.1). To ascertain that the adaptation score calculated over the data obtained during the MRI session was a stable measure of individual differences, the intraclass correlation(ICC) of the slopes across sessions 2, 3, 4 and 5 were calculated. Further, to evaluate the influence of contextual experimental effects (MRI vs lab-based assessments of pain habituation), ICCs were also calculated for sessions 3-5, which were all completed in the same lab. In the instance that ICCs indicated stability of adaptation slopes across sessions, average slopes were calculated, to evaluate correlations across baseline assessment variables, as well as the predictability of habitation in the MRI session for performance in the following behavioural sessions. All data were analysed using SPSS27(IBM Corp., Armonk, NY).

#### 2.4.2 MRI Data Analysis

##### 2.4.2.1. Pre-processing

FSL6.0[7] was used for all MRI analyses, including pre-processing, adhering to the protocol described by the CompCor[4]. Skull-stripping was performed using the Brain Extraction Tool (BET;[11]). Data were spatially smoothed using a 5mm FWHM Gaussian kernel, and interleaved slice timing correction was applied. To correct for B0 inhomogeneities in the data, the SE EPI acquisitions with opposite encoding directions were used to calculate a field map via TopUp[1] and applied to data. Functional data were registered to each subject’s anatomical space via Boundary-based registration (BBR) and registered to standard MNI space using 12-DOF non-linear transformation.

Using FAST[12], anatomical images were then segmented into grey matter, white matter (WM) and cerebrospinal fluid (CSF) masks. Masks were thresholded at .99, residuals were bandpass filtered (0.1/0.01Hz), normalised and time series for WM and CSF were generated. Using MCFLIRT[6], motion correction was applied, and this time-series was entered into a ‘nuisance removal’ GLM alongside WM and CSF time-series.

##### 2.4.2.2. Functional MRI Pain Task

The pre-processed single-subject data were modelled using pain stimulation events as explanatory variables, and then each of the four blocks of the pain task were concatenated using a fixed effects model as the second step. To capture changes in neural processes over time, alongside univariate concatenation (1,1,1,1), the data were also parametrically modulated in this second level analysis, to generate contrast maps for linear increases (-1.5, -0.5,0.5,1.5) or decreases (1.5,0.5, -0.5, -1.5) in activity across the blocks. All FEAT directories were entered into a third-level mixed effect FLAME1+2 analysis, wherein group mean, and pain adaptation slope (within the MRI task) were entered as explanatory variables. In line with considerations of inflated false-positive rates within parametric statistical methods[24], analyses were corrected for multiple comparisons using non-parametric permutation testing (Z>2.3, corrected p=.005), using the FSL Randomise toolbox[84]. Parameter estimates of significant clusters were extracted using FeatQuery. For post-hoc analyses, when large clusters encompassed multiple regions, probabilistic anatomical masks were thresholded at 50% probability, and used to extract values within the cluster specific to an anatomical region. Temporal dynamics in activity over the four runs were quantified by calculating the slope angle of parameter estimates over time. These neural slopes were then entered into separate linear regression models to predict single-session temporal summation scores as well as pain habituation slopes calculated across the ratings obtained in sessions 3, 4 and 5.

##### 2.4.2.3. Resting-state MRI

In order to examine the connectivity patterns that might support regions associated with individual differences in adaptation to pain, clusters originating from the analysis of the functional pain task were used as ROIs in a resting state PPI analysis. Using the same cluster masking from parameter extraction in the previous analysis, larger clusters which encompass multiple regions were masked, to facilitate conclusions of anatomical specificity. Psychophysiological interactions were investigated by extracting mean time series for regions-of-interest (significant clusters from the fMRI pain task), which were included as predictors in a whole-brain connectivity analysis. Contrast maps were then entered into a higher-level analysis with demeaned adaptation slopes, derived from MRI task pain ratings, as between-subjects regressors. This resulted in regions whose connectivity with the seed region was significantly correlated with pain adaptation. Analyses were corrected for multiple comparisons using Gaussian random field theory (z>2.3, p<.05). Using FeatQuery, parameter estimates were extracted from clusters showing individual differences in functional connectivity with each ROI as a function of adaptation slopes. To restrict the extracted data to grey matter regions, each cluster was multiplied using masks within the Harvard-Oxford anatomical atlases, using a 25% probabilistic threshold.

## 3.1 Results

### 3.1.1 Pain Adaptation

Across the sample, 37.6% (n=32) of participants demonstrated overall habituation to painful stimulation during the MRI task, as opposed to 57.6%(n=49) who sensitized, and 4.7% (n=4) who showed no change. Pain rating slope were stable across all four sessions (ICC(2,1)=.47(.36-.58), p<.001; Fig.2). The data indicate the presence of context effects on reported pain over time, represented by higher stability when excluding the MRI session (ICC(2,1)=.64(.57-.74),p<.001). As the slopes across sessions 3-5 were shown to be stable, a single average slope value (average rate of change) was calculated and correlated with relevant variables obtained during the baseline sessions. Pain adaptation during the MRI task significantly predicted the average slope in the following sessions (F(1,83)=7.86,p=.006, R^2^=.086). Regarding pain adaptation, those with higher predisposition to sensitization showed higher temporal summation (r(85)=.41,p<.0001), neuroticism (r(84)=.37,p<.001) and depression scores (r(85)=.27,p<.05), as well as lower trait mindfulness (r(85)=-.26,p<.05).

**Figure 2.**
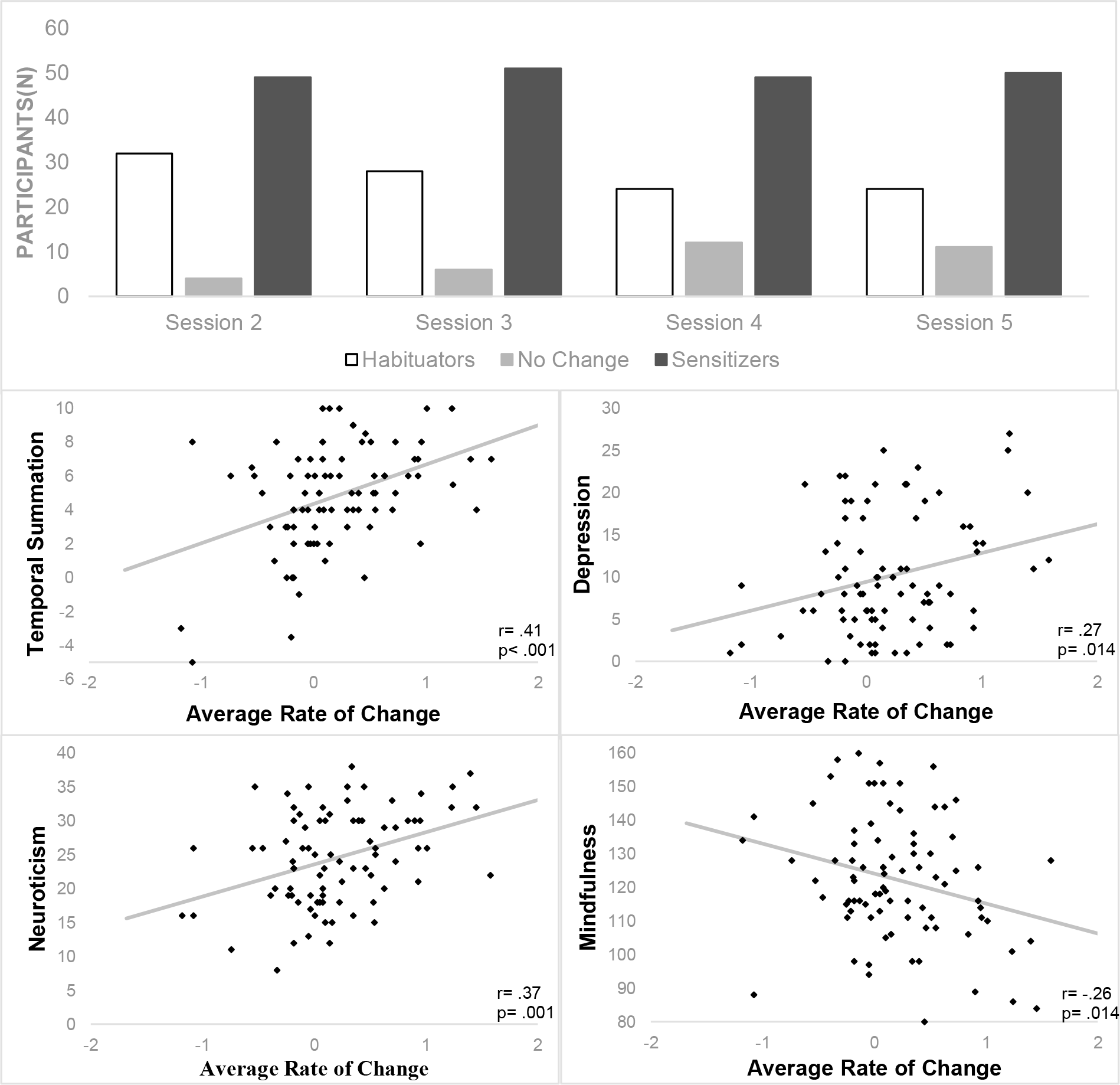
(top) Number of participants within each session that habituated, sensitized, or showed no change to repetitive painful stimulation. (bottom) Correlations between average pain adaptation slope across all four sessions (higher value; predisposition to sensitization), and baseline psychobehavioural variables.

### 3.1.2 fMRI Pain Task

#### 3.1.2.1. Main Effects of Pain Stimulation

Pain stimulation activated brain areas frequently associated with pain (Supp.1), including bilateral insula, secondary somatosensory, premotor, and cingulate cortices, as well as the thalamus, bilateral amygdala, and brainstem. Parametric modulation analysis identified regions that demonstrated significant linear increases or decreases in activation during pain stimulation over the four blocks. Significant increases in activation were identified in the primary somatosensory and motor cortices, as well as in the parietal lobe and V1. Although, significant decreases in activation were identified in the right secondary somatosensory cortices, extending into the parietal operculum and Broca’s area, these did not survive correction following non-parametric permutation testing.

#### 3.1.2.2. Dynamic Mechanisms of Pain Adaptation

Pain adaptation was associated with activity in the right anterior hippocampus and amygdala and right premotor cortex, bilateral primary somatosensory, and motor cortices across the four blocks (Fig. 3). Such that, habituation was associated with increasing activity in the hippocampus and amygdala, and decreased activity in sensorimotor regions. Parameter estimates were extracted from each cluster, and masked using probabilistic atlases (amygdala, hippocampus, somatosensory cortices and motor cortices). Activity change in the hippocampus over the four runs significantly predicted pain adaptation in the future behavioural sessions (F(1,83)=4.03,p<.05, R^2^=.05) and temporal summation scores(F(1,83)=4.72,p<.05).

**Figure 3.**
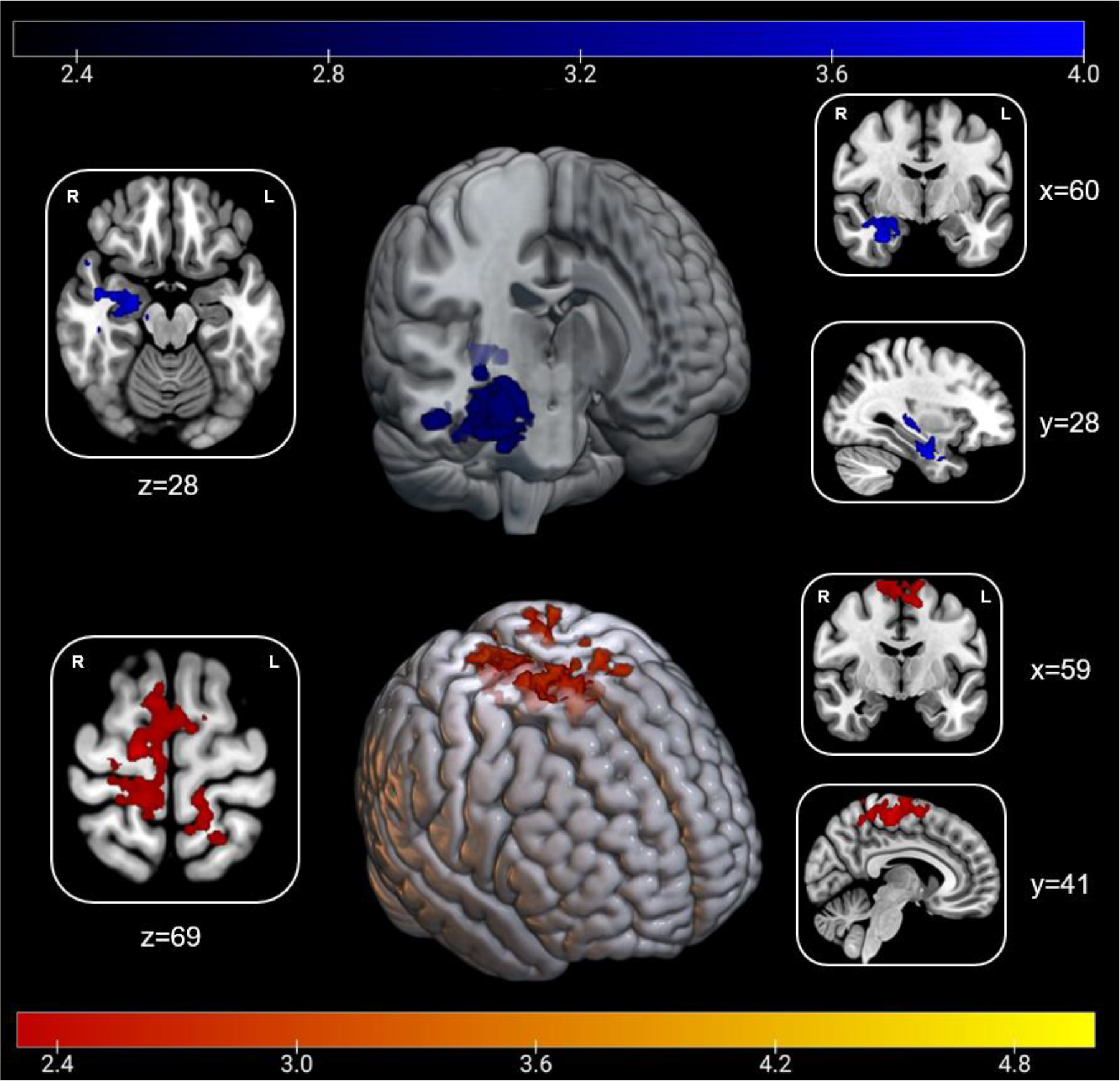
Dynamic mechanisms of pain adaptation |(top) Cluster encompassing the right anterior hippocampus and amygdala (Z_max_=3.97; x=28,y=-6,z=-30) where increasing activity is associated with habituation to pain (bottom) Cluster encompassing the right premotor cortex and bilateral somatosensory and motor cortices (Z_max_=4.49; x=6,y=-20,z=66) where increasing activity is associated with sensitization. Numbers denote slice depicted.

### 3.1.3. Resting-state MRI

#### Seed-based Whole-brain Functional Connectivity

Functional connectivity analyses revealed pain adaptation was associated with variations in connectivity between the right anterior hippocampus and ventromedial prefrontal cortex at rest, such that individuals with higher rates of habituation had higher connectivity (Fig. 4). Habituation was also associated with higher functional connectivity between the sensorimotor cluster from the event-related analysis and a cluster extending across the right hippocampus, parahippocampal gyrus, amygdala, and insula cortex. This cluster partially overlapped with the cluster derived from the event-related habituation analysis. Resting-state analysis therefore yielded two connectivity pairings: Firstly, the hippocampus seed to ventromedial prefrontal cortex and, second, the sensorimotor seed to hippocampus.

**Figure 4.**
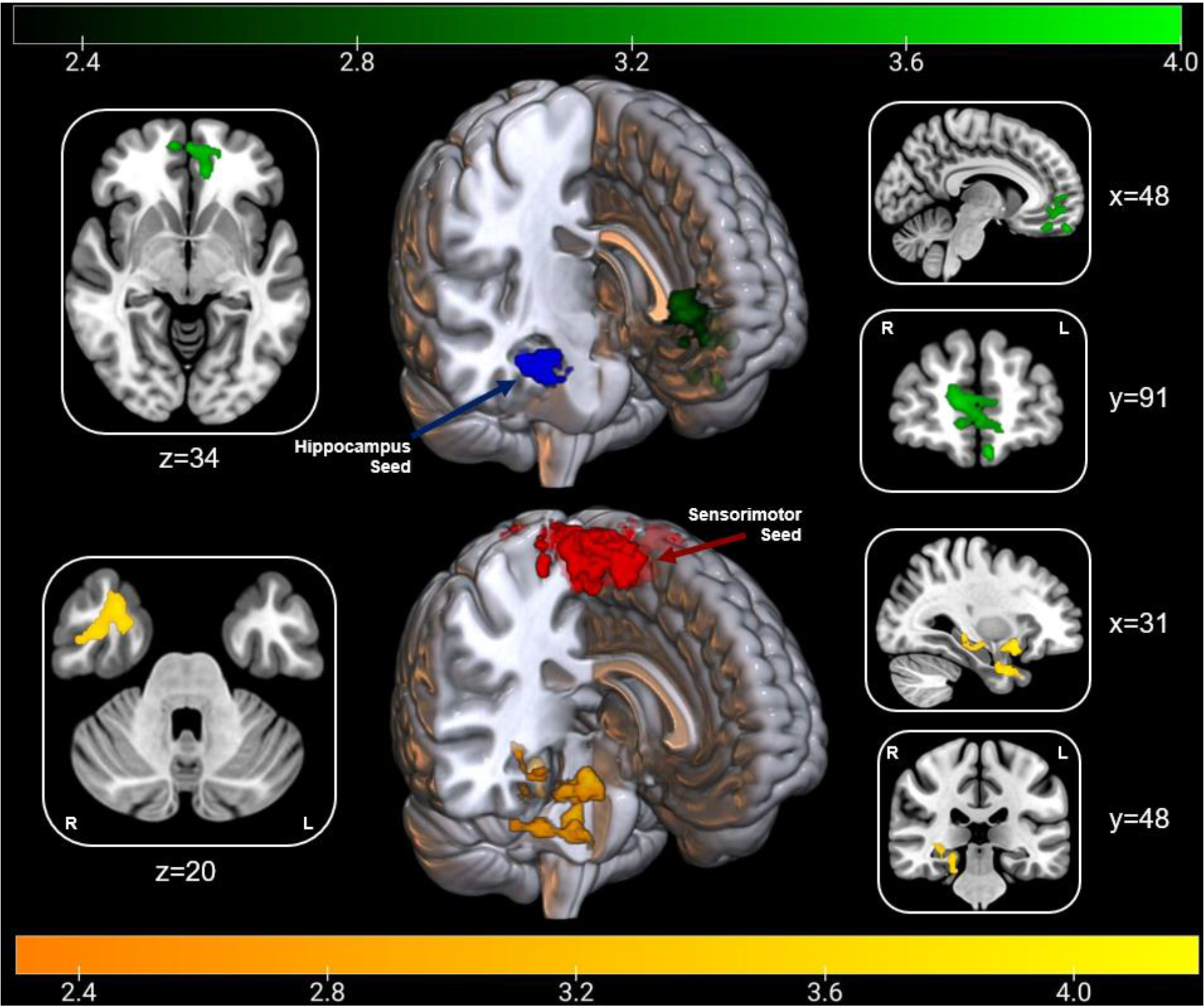
Functional Connectivity of Habituation |(top) Seed-based functional connectivity of the right hippocampus seed(blue) where habituation was associated with higher connectivity to the ventromedial prefrontal cortex(green; Z_max_=3.78; x=-6,y=58,z=-22) |(bottom) Seed-based functional connectivity of the sensorimotor seed(red) where habituation was associated with higher connectivity to the right hippocampus, amygdala, and insula cortex(orange; Z_max_=4.3; x=30,y=12,z=-12). Numbers denote slice depicted.

## Discussion

This study investigated the stability of individual differences in pain adaptation, and their underlying neural mechanisms. We observed substantial individual differences in response to repetitive nociceptive stimulation. Importantly, these individual patterns of pain adaptation were reliable across sessions, and experimental contexts. Pain adaptation was correlated with temporal summation, such that sensitization was associated with greater summation. Adaptation was also correlated with psychological variables, with higher levels of sensitization associated with higher depression and neuroticism, and lower trait mindfulness. Higher levels of habituation during repetitive nociceptive stimulation, were associated with increasing activity in the anterior hippocampus and amygdala over time, whereas sensitization was associated with increasing activity in the somatosensory, motor and premotor cortices. Activity change within the hippocampus in response to repeated nociceptive stimulation predicted pain adaptation in future behavioural sessions, as well as single-session temporal summation scores. In the resting state functional connectivity analysis, habituation was associated with higher connectivity between the hippocampus/amygdala seed region and vmPFC, and higher connectivity between the sensorimotor cluster and the hippocampus, amygdala, and insula cortex. These findings demonstrate that pain adaptation involves a network of brain regions typically associated with the interpretation and evaluation of pain and is not solely a product of variation in nociceptive processing. As such, the way in which an individual processes pain is an important determinant of sensitization and habituation and, importantly, is a stable phenotypic quality.

How individuals respond to repetitive nociceptive stimulation is important to understanding vulnerability to pain in both experimental and clinical environments. In circumstances where heightened risk is perceived, pain responses will be enhanced (sensitization). If unavoidable noxious stimuli are perceived to pose no threat, pain habituation may help preserve resources. When empirically examining group effects, proportions of habituators and sensitizers reported varies greatly[19,25,70,27,29,32,37,45,50,51,68]. Our finding is the first to indicate that the propensity for an individual to habituate or sensitize to pain is stable within individuals across time and context and, hence, may serve as a valuable phenotypic marker.

### Sensory-discriminative habituation

Pain adaptation was associated with neural activity in the premotor, motor, and somatosensory cortices, such that habituation was associated with reduced activity over time. Reduced sensory-discriminative processing facilitating lower pain is well documented[33,41] and this process appears to be disrupted in chronic pain patients[17,43]. Using a similar design, it has been reported that stimulation over 8 separate days was associated with reduced BOLD responses to nociceptive stimuli in SII, as well as in the insula, thalamus and putamen[8]. Those data also indicated that pain adaptation is associated with increased grey matter density in the somatosensory cortex[77], which was maintained at 21-day follow-up. Importantly, most studies quantified habituation/sensitization exclusively by examining brain activity during nociceptive stimulation, without including pain ratings. Consequently, the neural results are predominantly focused on sensory-discriminative regions and are likely tracking an automatic habituatory effect via reduced nociceptive processing. This current study advances our understanding of the subjective experience of pain adaptation by focusing on participant pain ratings, thus focusing on the phenomenological experience of pain, alongside the sensory dimension of nociception.

### Hippocampal-Amygdala Circuitry

Activity in the anterior hippocampus and amygdala was also associated with pain adaptation, with habituation linked to increasing activity over time. In combination, the interaction of amygdala-hippocampal processes are associated with the processing of emotional events and representations[65,72]. The anterior hippocampus is associated with pain information processing[39], mediation of context-dependent pain expectancy[89] and integrating multiple sources of information to make decisions[87]. Chronic pain is associated with reduced bilateral hippocampal volume[88], disrupted hippocampal neurogenesis[20,42] and increased neuroinflammation[79]. Mirroring our findings, coupling between the hippocampus and mPFC has been identified as a biomarker for the transition from acute to chronic pain[44]. The strength of this connectivity can predict variations in back pain across the year and the likelihood of recovery from subacute pain, with persistent back pain associated with large decreases in prefrontal-hippocampal functional connectivity. Our finding furthers this model by highlighting that vmPFC-hippocampus connectivity in healthy controls facilitates pain modulation, which may provide clarification to the mechanisms that predispose certain individuals to chronic pain. The underlying behavioural function of this biomarker could be pain adaptation, with those more likely to habituate to pain being best positioned for the resolution of pain before it develops into a persistent chronic condition. The expense of MRI often prevents its clinical use in pain assessment, making a behavioural proxy for a biomarker an asset. Whether our pain adaptation protocol could be applied to predict this transition from acute to chronic pain requires further investigation.

The amygdala is also an important contributor to the emotional component of pain[28]. The latero-capsular division of the central nucleus of the amygdala, or “nociceptive amygdala”, is associated with the integration of external and internal environmental information with nociception[49,83]. Intuitively, the direction of this change in neural activity is surprising, given the relationship between the amygdala and pain intensity[11] and depressive or anxious states[3,18]. However, the amygdala is also involved in regulating moment-by-moment vigilance levels[81] and the processing of emotionally salient stimuli, especially when pertinent for later evaluation[15]. Within our design, participants were not informed their stimuli were all the same temperature and were asked to rate their average pain intensity at the end of a trial. As pain involves an emotional component[66], our results suggest that the hippocampus and amygdala facilitate the consolidation and appraisal of pain across repetitive stimulation via emotional appraisal processes. This mechanism may have facilitated accurate appraisal that stimulation intensity was not increasing or risking physical damage, thus reducing threat appraisal and, ultimately, facilitating a beneficial habituatory response.

### Pain Adaptation as a Phenotype

The influence of pain sensitization on pain chronicity is an crucial area of study[5,52], focusing on the discrepancy between peripheral drivers of pain, and the perceptual consequences. Pro-nociceptive phenotypes have long been investigated to identify those at risk of chronification of pain or poor treatment response[85]. Psychometrics, such as the central sensitization inventory(CSI), may be used for this purpose[48], although it is unclear if CSI actually quantifies its namesake construct[2]. Instead, QST is a more appropriate tool for quantifying these profiles[1,12,34], with temporal summation frequently cited as a predictive tool for pain chronification and poor clinical outcomes[6,61,80]. Indeed, in our data, TS was correlated with adaptation slopes and hippocampal activity dynamics. Taken together, pain adaptation may be a useful variable for quantifying nociceptive phenotypes.

Habituation may represent an individual appropriately evaluating their environment as posing no risk of harm, facilitating the reduction of needlessly expended defensive resources. Sensitization to pain elicits the opposite and poses a risk of cognitive drain and increased pain sensitivity. Our data indicate that psychosocial features influence an individual’s predisposition to either side of this continuum. Neuroticism is a trait commonly associated with exaggerated threat-appraisal, and sensitivity to environmental stress[82]. Neuroticism is consistently associated with the transition from acute to chronic pain[60], the development of chronic post-surgical pain[26], impaired quality of life[67] and chronic pain sufferers frequently report higher neuroticism scores than healthy control[35]. Relatedly, depression is highly co-morbid with pain[69] and is associated with poor clinical outcomes[53,63], and pain-related interference and disability[38]. Within our data, both neuroticism and depression were correlated with pain adaptation, with higher scores facilitating pain sensitization. Conversely, mindfulness, which was associated with pain habituation in our data, is characterised as a non-judgemental present-moment attentional regulation[10], associated with lower pain sensitivity[31] and higher functioning in chronic pain patient[40]. Our findings underscore that pain adaptation is a volitional process, rather than solely an involuntary habituation in nociceptive processing. Functional connectivity analyses at rest further indicate that participants with decreasing sensorimotor activation during the task, who habituated to pain, showed increased connectivity with regions of the brain associated with emotional responding and learning (hippocampus, amygdala and insula). In turn, these regions were increasingly coupled with the vmPFC, a key region for emotional regulation, via flexible value assignment and the inhibition of activity in the amygdala depending on context[4,71]. This, not only, expands on the pain adaptation phenotype, but also underscores the importance of emotional regulation in pain resilience and, by extension, psychological interventions to correct a pro-nociceptive phenotype.

### Summary

This study is the first to evidence the stability of individual differences in pain adaptation and underscore its utility as a phenotypical variable, potentially suitable for clinical use. Pain adaptation measured in a single session can predict similar rates of change in subsequent sessions, and is associated with temporal summation, depression, neuroticism and mindfulness. Our findings suggest that reduced sensory-discriminative and increased amygdala-hippocampal activity, alongside interaction with the insula and vmPFC facilitates habituation. It is likely that adaptation involves components of, both, automatic and volitional processes, which include the appraisal of pain and the regulation of emotion. Future work is needed to explore if these neural mechanisms represent biological markers for vulnerability to prolonged nociceptive input or whether a repetitive pain stimulation paradigm can be used as a phenotypical assessment to predict pain chronicity.

## Contributions that do not justify authorship

None

## Technical assistance

The Centre for Integrative Neuroscience & Neurodynamics Operational Support Team at the University of Reading

## Financial and material support

RH:-Medical Research Council MR/R005656/1

TS:-Medical Research Council MR/R005656/1

WG:-Leverhulme Early Career Fellowship

GA:-Medical Research Council MR/R005656/1

## Financial support that may represent a conflict of interest

None

## Open Access Statement

For the purpose of open access, the author has applied a Creative Commons Attribution (CC BY) licence to any arising Author Accepted Manuscript version

