## Supplemental Figure 1 for "Temporal dynamics of hippocampal activity predict stable patterns of sensitization or habituation to noxious stimulation across sessions"

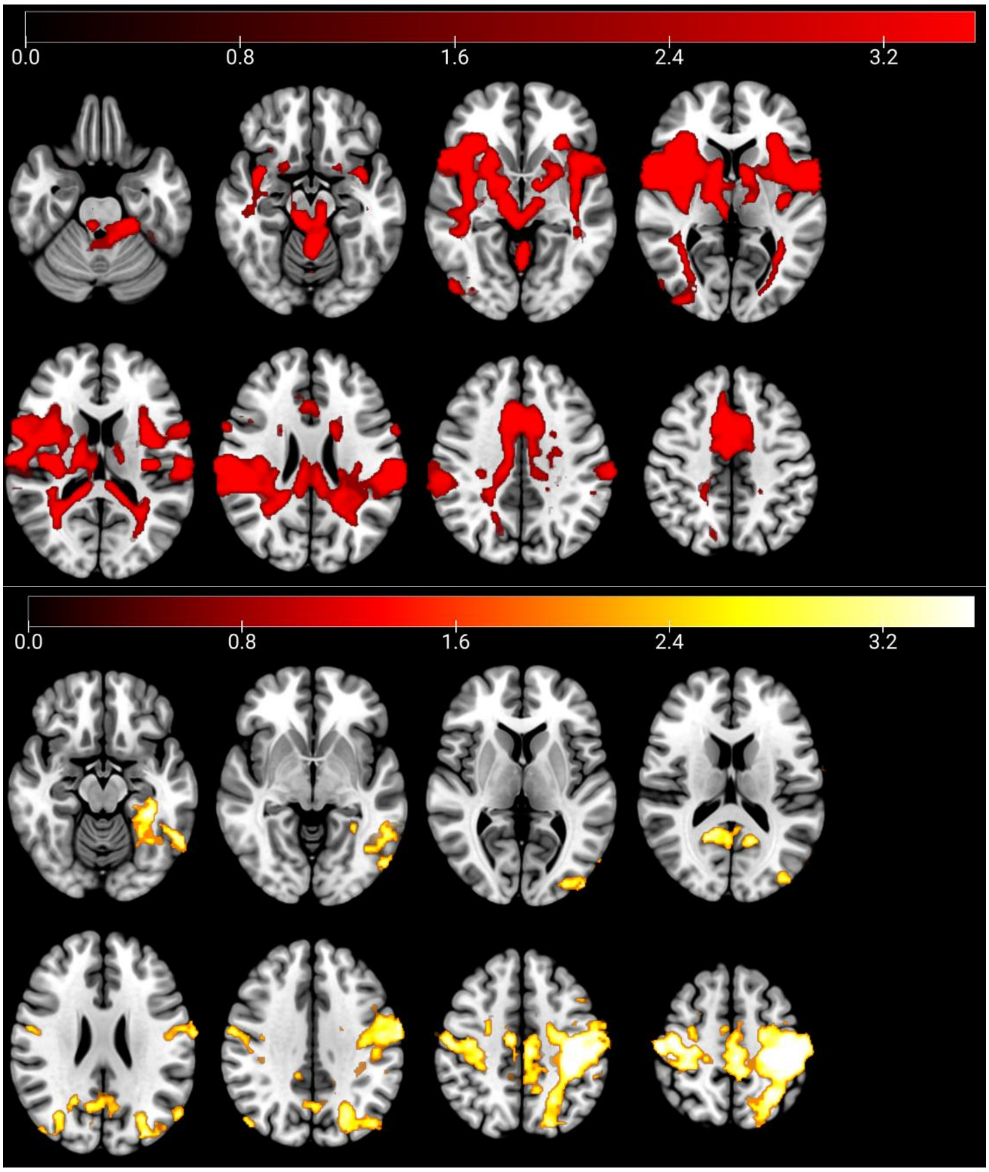


**Supplementary 1a** | Main effect of pain stimulation (top) encompassing bilateral amygdala, insula, somatosensory, premotor and cingulate cortices, thalamus and brainstem ($Z_{max}$=3.54,x=54,y=-38,z=20). Dynamic increases in activity over time were identified via parametric modulation analyses (middle) in the somatosensory and motor cortices, parietal lobe and V1 ($Z_{max}$=3.54,x=32,y=-32,z=48).
